# Genetic architecture of individual meiotic crossover rate and distribution in a large Atlantic Salmon (*Salmo salar)* breeding population

**DOI:** 10.1101/2023.06.07.543993

**Authors:** Cathrine Brekke, Susan E. Johnston, Tim M. Knutsen, Peer Berg

## Abstract

Meiotic recombination through chromosomal crossovers ensures proper segregation of homologous chromosomes in meiosis, while also breaking down linkage disequilibrium and shuffling alleles at loci located on the same chromosome. Rates of recombination can vary between species, but also between and within individuals, sex and chromosomes within species. Indeed, the Atlantic salmon genome is known to have clear sex differences in recombination with female biased heterochiasmy and markedly different landscapes of crossovers between males and females. In male meiosis, crossovers occur strictly in the telomeric regions, whereas in female meiosis crossovers tend to occur closer to the centromeres. However, little is known about the genetic control of these patterns and how this differs at the individual level. Here, we investigate genetic variation in individual measures of recombination in >5000 large full-sib families of a Norwegian Atlantic salmon breeding population with high-density SNP genotypes. We show that females had 1.6× higher crossover counts (CC) than males, with autosomal linkage maps spanning a total of 2174 cM in females and 1483 cM in males. However, because of the extreme telomeric bias of male crossovers, female recombination is much more important for generation of new haplotypes with 8x higher intra-chromosomal genetic shuffling than males. CC was heritable in females (h^2^ = 0.11) and males (h^2^ = 0.10), and shuffling was also heritable in both sex but with a lower heritability in females (h^2^ = 0.06) than in males (h^2^ = 0.11). Inter-sex genetic correlations for both traits were close to zero, suggesting that rates and distribution of crossovers are genetically distinct traits in males and females, and that there is a potential for independent genetic change in both sexes in the Atlantic Salmon. Together, these findings give novel insights into the genetic architecture of recombination in salmonids and contribute to a better understanding of how rates and distribution of recombination may evolve in eukaryotes more broadly.

## Introduction

Meiotic recombination is a fundamental part of sexual reproduction, where chromosomal crossing-over between the maternal and paternal chromosomes during synapsis in the early prophase of meiosis, leads to novel combinations of alleles in the gametes transmitted to the next generation. It is of large interest in studies of both wild and domestic species, as it breaks up linkage between loci located on the same chromosome and in turn affects the speed and degree of responses to selection (1,2). Recombination also has a mechanistic role in the proper segregation of chromosomes during meiosis; a lack of crossovers can lead to harmful outcomes, such as aneuploidy (i.e. the incorrect number of chromosomes in gametes); (3–6), whereas high rates of recombination can be associated with increased mutation rates at crossover sites (7). Yet, there is large variation in recombination rates within and between species, populations, sexes, individuals, and chromosomes across sexually reproducing eukaryotes (8,9), suggesting a combination of mechanistic and evolutionary processes in driving this variation.

Studies of individual crossover counts in mammals and birds often show that it is variable and heritable, with a conserved, oligogenic architecture (10–18). Genes including *RNF212*, *REC8, SPO11* and *RNF212B* are repeatedly associated with recombination rate in mammal and bird studies (19–24) whereas the locus *PRDM9* has been identified as a gene that determines recombination hotspot positioning in mammals (25). Furthermore, most species studied show sex differences in the degree and direction of recombination rates and landscapes (known as heterochiasmy) and underlying genetic architectures often differ between the sexes, through sex-limited or sex-differential effects of associated loci (9,12–17,21,26,27)). The causes and consequences of this sexual dimorphism has been of interest for decades, with arguments often centring around sex differences in the fitness consequences of preserving beneficial linked alleles (“haplotypes”) or generating novel haplotypes that increase gamete and/or offspring fitness (28–31). However, understanding the broader scale evolution of heterochiasmy remains to be fully understood, due to a lack of suitable empirical data to test hypotheses (30,32–36). Notably, these previous comparative studies have mainly focussed on differences in linkage map lengths and/or crossover counts, yet the evolutionary arguments above centre around the preservation/generation of haplotypic variation. However, theoretical work has shown that the rate of allelic shuffling (i.e. the uncoupling of linked allelic variation on chromosomes) is influenced by crossover positioning, where a crossover on the middle of a chromosome will lead to higher rates of allelic shuffling than a crossover on distal regions (37). Therefore, studies of heterochiasmy should consider not only sex-differences in crossover counts, but also in rates of allelic shuffling within chromosomes, to better disentangle processes related to chromosome disjunction and genetic linkage.

The Salmonidae family (salmon, char, trout, whitefish and grayling) share an ancestor that underwent a whole genome duplication (WGD) event some 50-100 million years ago (38). Studies in salmonids find that chromosome arms that still show high sequence similarity exchange genetic material during meiosis in a quadrivalent formation (39), which appears to be almost exclusive to male meiosis (40,41), and therefore may lead to different recombination patterns between the sexes. Indeed, studies in Atlantic salmon (*Salmo salar*) have shown extreme and distinct differences in recombination landscapes between males and females (42,43). An early linkage analysis in Atlantic salmon with 442 markers showed little to no recombination in male linkage groups, implying that female recombination was 5 times higher than that of males (44). However, as marker densities increased (to 5650 markers), it was discovered that males do recombine with an overall rate relatively close to females in most chromosomes, but almost exclusively in the telomeric regions (42), a pattern which could not be picked up by lower-density arrays. These stark differences in landscape indicate the presence of large sex-differences in allelic shuffling between linked loci in these species, but the causes and consequences of variation in meiotic crossovers remain unknown. In this study, we investigated individual-, sex- and population-level variation in meiotic crossover rates and landscapes in a large breeding population of Atlantic salmon. We use data from more than 5000 full sib families with genotypes on ∼35,000 SNP markers to: (a) construct sex-specific linkage maps; (b) quantify individual crossover counts and rates of intra-chromosomal allelic shuffling; (c) determine their heritabilities and genetic architectures within each sex; and (d) investigate cross-sex genetic correlations.

## Results

### Linkage mapping

The sex-specific linkage maps spanned a total of 2173.80 cM in females and 1482.96 cM in males, with the female map 1.47 times than in males (Table 1, Fig 1). In agreement with Lien et al. (2011), the biggest sex differences were on chromosomes 2, 8 and 17, where the female maps were 5.34, 28.13 and 16.09 times longer than the male maps, respectively. In these three chromosomes, we detected very few crossover events in males, with male maps only 2-20 cM long. Obligate crossing-over results in a minimum predicted map length of 50cM; therefore, we assume that we are unable to pick up all crossovers in males on their chromosomes, perhaps either due to lower marker coverage in the telomeric regions on these chromosomes or due to multivalent pairing and crossing over with a different pair of chromosomes (39). For the remaining chromosomes, the genetic map length is 1 to 2.1 times longer in females. Chromosome-level results from the linkage mapping for all chromosomes are provided in Table 1, and Marey maps showing the relationship between the physical and genetic length of the chromosomes are shown in Fig 1. The full linkage map can be found in

**Fig 1.**
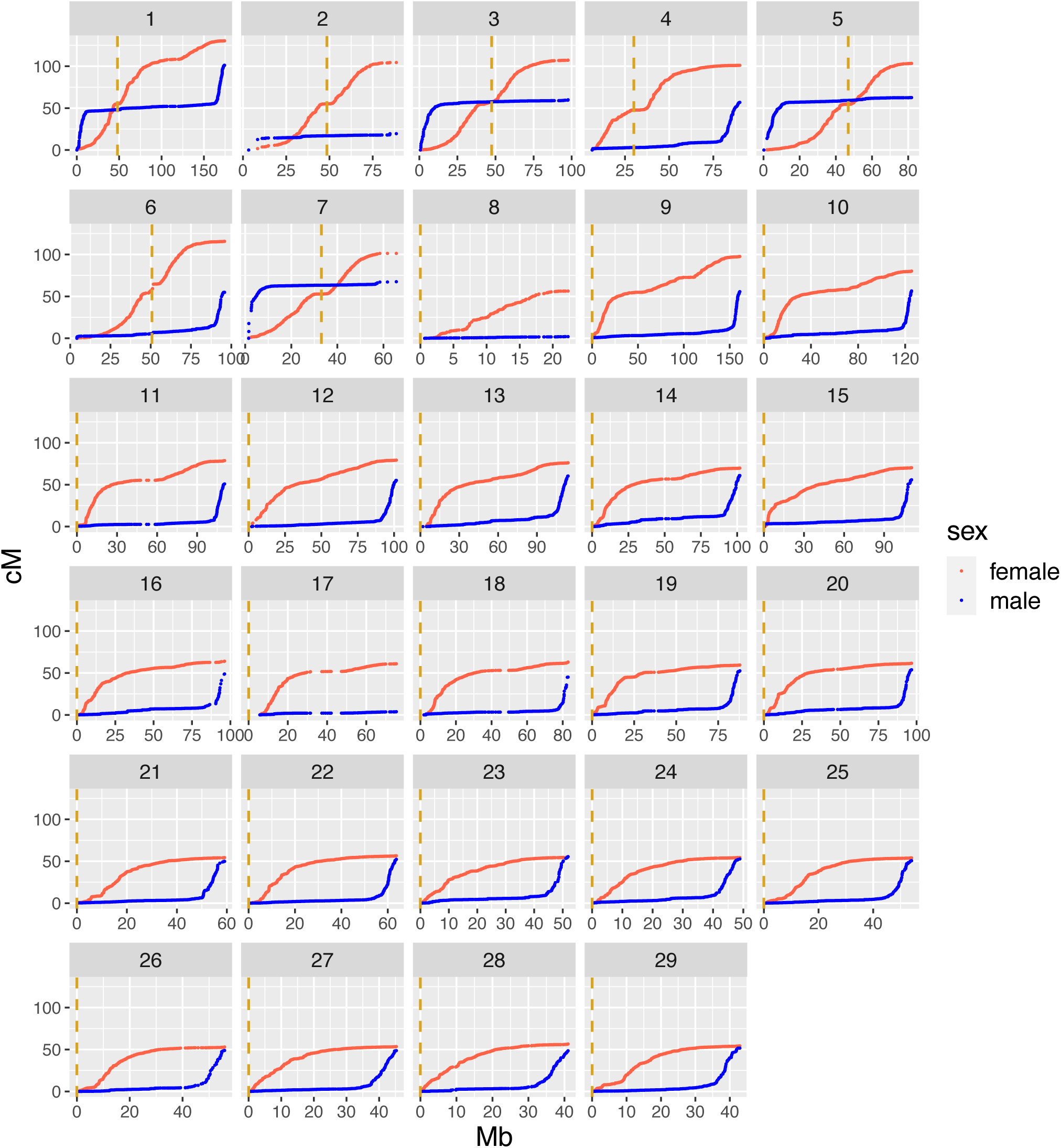
Male and female Marey maps for the 29 Atlantic salmon chromosomes. The physical position in Mb is plotted against the genetic position in cM for the SNP markers within each linkage group. Female positions in red and male in blue. The dashed vertical lines in yellow are the centromere positions from the reference genome Ssal_v3.1 (GenBank accession GCA_905237065.1).

**Table 1.** Summary of linkage mapping results by sex and chromosome.

| Chr | Number of markers | Chr length (Mb) | Male cM | Female cM | Female/male ratio | Male rate (cM/Mb) | Female rate (cM/Mb) |
| --- | --- | --- | --- | --- | --- | --- | --- |
| ssa1 | 2828 | 174.50 | 101.15 | 130.32 | 1.29 | 0.58 | 0.75 |
| ssa2 | 749 | 95.48 | 19.54 | 104.45 | 5.34 | 0.20 | 1.09 |
| ssa3 | 1417 | 105.78 | 59.92 | 107.26 | 1.79 | 0.57 | 1.01 |
| ssa4 | 1485 | 90.54 | 56.88 | 101.11 | 1.78 | 0.63 | 1.12 |
| ssa5 | 1246 | 92.79 | 62.60 | 103.35 | 1.65 | 0.67 | 1.11 |
| ssa6 | 1255 | 96.06 | 54.88 | 115.71 | 2.11 | 0.57 | 1.20 |
| ssa7 | 852 | 68.86 | 67.55 | 101.37 | 1.50 | 0.98 | 1.47 |
| ssa8 | 300 | 28.86 | 2.01 | 56.40 | 28.13 | 0.07 | 1.95 |
| ssa9 | 2333 | 161.28 | 55.68 | 97.65 | 1.75 | 0.35 | 0.61 |
| ssa10 | 1881 | 125.88 | 56.54 | 80.23 | 1.42 | 0.45 | 0.64 |
| ssa11 | 1392 | 111.87 | 50.92 | 78.75 | 1.55 | 0.46 | 0.70 |
| ssa12 | 1485 | 101.68 | 55.15 | 79.27 | 1.44 | 0.54 | 0.78 |
| ssa13 | 1611 | 114.42 | 60.33 | 76.21 | 1.26 | 0.53 | 0.67 |
| ssa14 | 1557 | 101.98 | 60.94 | 69.79 | 1.15 | 0.60 | 0.68 |
| ssa15 | 1524 | 110.67 | 55.84 | 70.27 | 1.26 | 0.50 | 0.63 |
| ssa16 | 1302 | 96.49 | 48.83 | 64.02 | 1.31 | 0.51 | 0.66 |
| ssa17 | 681 | 87.49 | 3.79 | 60.96 | 16.09 | 0.04 | 0.70 |
| ssa18 | 1067 | 84.08 | 45.04 | 63.04 | 1.40 | 0.54 | 0.75 |
| ssa19 | 1157 | 88.11 | 52.57 | 59.48 | 1.13 | 0.60 | 0.68 |
| ssa20 | 1331 | 96.85 | 53.81 | 61.61 | 1.14 | 0.56 | 0.64 |
| ssa21 | 962 | 59.82 | 49.88 | 54.37 | 1.09 | 0.83 | 0.91 |
| ssa22 | 1122 | 63.82 | 52.31 | 56.56 | 1.08 | 0.82 | 0.89 |
| ssa23 | 949 | 52.46 | 55.52 | 55.48 | 1.00 | 1.06 | 1.06 |
| ssa24 | 925 | 49.35 | 52.75 | 54.53 | 1.03 | 1.07 | 1.10 |
| ssa25 | 888 | 54.39 | 50.67 | 54.06 | 1.07 | 0.93 | 0.99 |
| ssa26 | 593 | 55.99 | 48.91 | 53.05 | 1.08 | 0.87 | 0.95 |
| ssa27 | 825 | 45.31 | 48.75 | 53.57 | 1.10 | 1.08 | 1.18 |
| ssa28 | 606 | 41.47 | 48.55 | 56.60 | 1.17 | 1.17 | 1.36 |
| ssa29 | 709 | 43.05 | 51.66 | 54.34 | 1.05 | 1.20 | 1.26 |
| Total | 35032 | 2499.33 | 1482.96 | 2173.80 | 1.47 | 0.59 | 0.87 |

Mb is the physical length of the chromosomes in megabases. cM is the estimated genetic length of the chromosomes in centiMorgan. Male and female rate is the recombination rate in cM/Mb. Total is the the total lengths and rates of the 29 chromosomes. Physical length of the chromosomes are from reference genome Ssal\_v3.1(GenBank accession GCA\_905237065.1)

### Fine-scale recombination rates along the genome

Patterns of recombination rates across the genome were strikingly different between males and females, in agreement with previous studies (42,43). Male recombination rates were highly elevated in the sub-telomeric regions (e.g. up to ∼10Mb from telomeres) and almost non-existent in the rest of the genome (Fig 2). Conversely, female recombination rates were elevated in peri-centromeric regions and reduced in telomeric regions (Fig 2). In immediate proximity to the centromere, both male and female recombination rates are close to zero in most chromosomes, indicating that the centromere itself has a suppressive effect on recombination (Fig 2).

**Fig 2.**
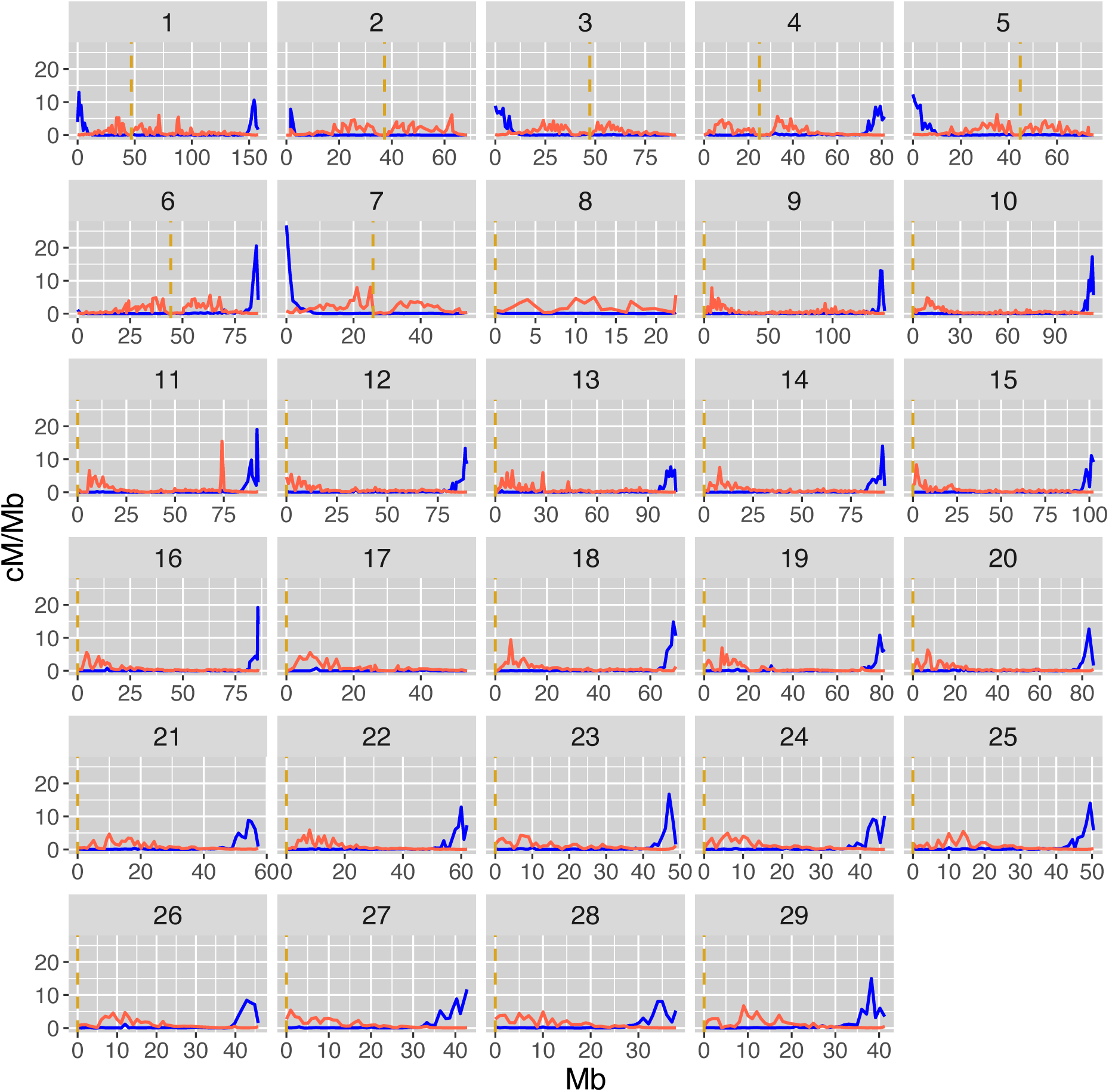
Fine-scale sex-specific recombination rate along the 29 Atlantic salmon chromosomes. The recombination rate in cM/Mb within each 1Mb bin for males in blue and females in red. The dashed vertical lines in yellow are the centromere positions from the reference genome Ssal_v3.1 (GenBank accession GCA_905237065.1).

### Individual recombination rates

The total numbers of observations (i.e. individual gametes) were 287,063 from 1,580 unique females, and 287,111 from 889 unique males. Crossover count (CC) was approximately normally distributed in both males and females (Fig 3), with a mean of 19.6 (± 3.7 SD) in females and 12.1 (± 2.9 SD) in males (Table 2). Intra-chromosomal genetic shuffling (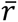) was normally distributed in males and females (Fig 4), with a mean of 8.06×10^-3^ (± 1.87×10^-3^ SD) in females and 1.01×10^-3^ (± 0.56×10^-3^ SD) in males, representing 8-fold higher levels of shuffling in females (Table 2).

**Fig 3.**
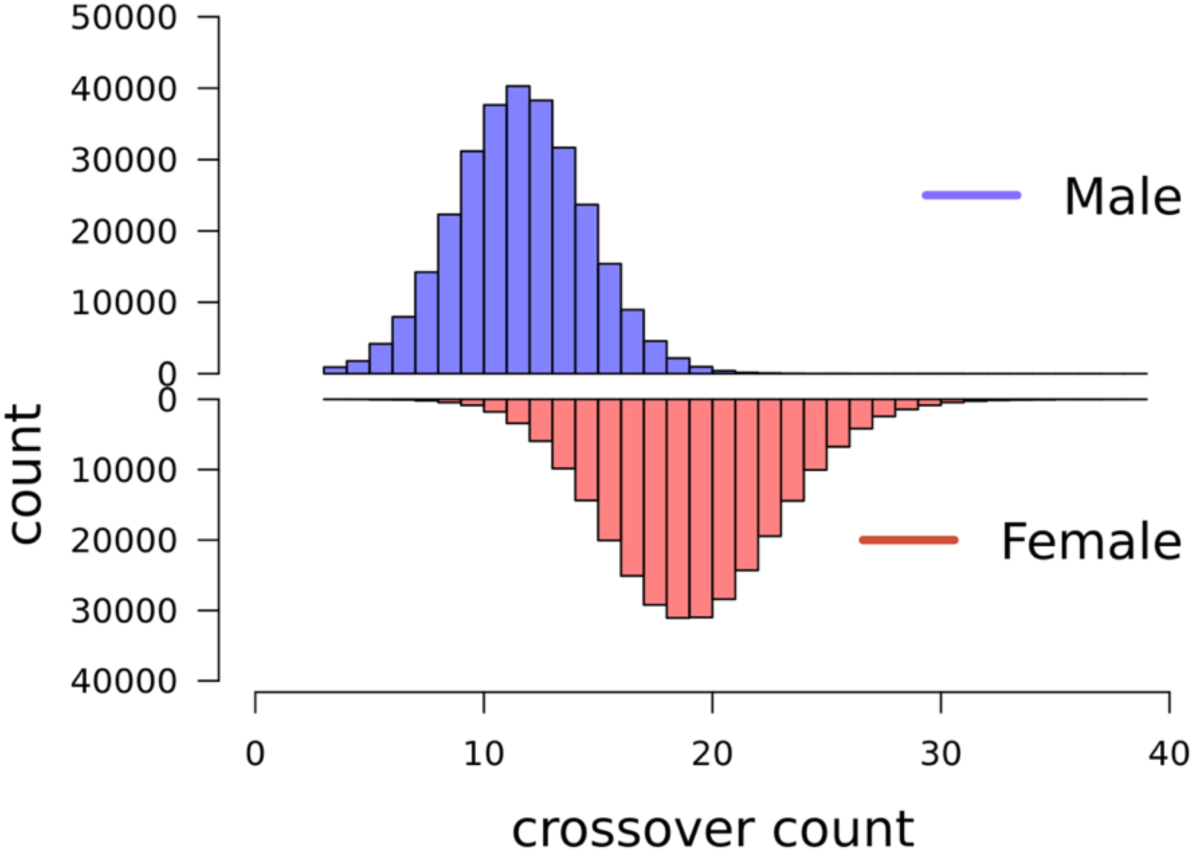
Distribution of crossover count (CC). for paternal gametes in blue (top) and maternal gametes in red (bottom).

**Fig 4.**
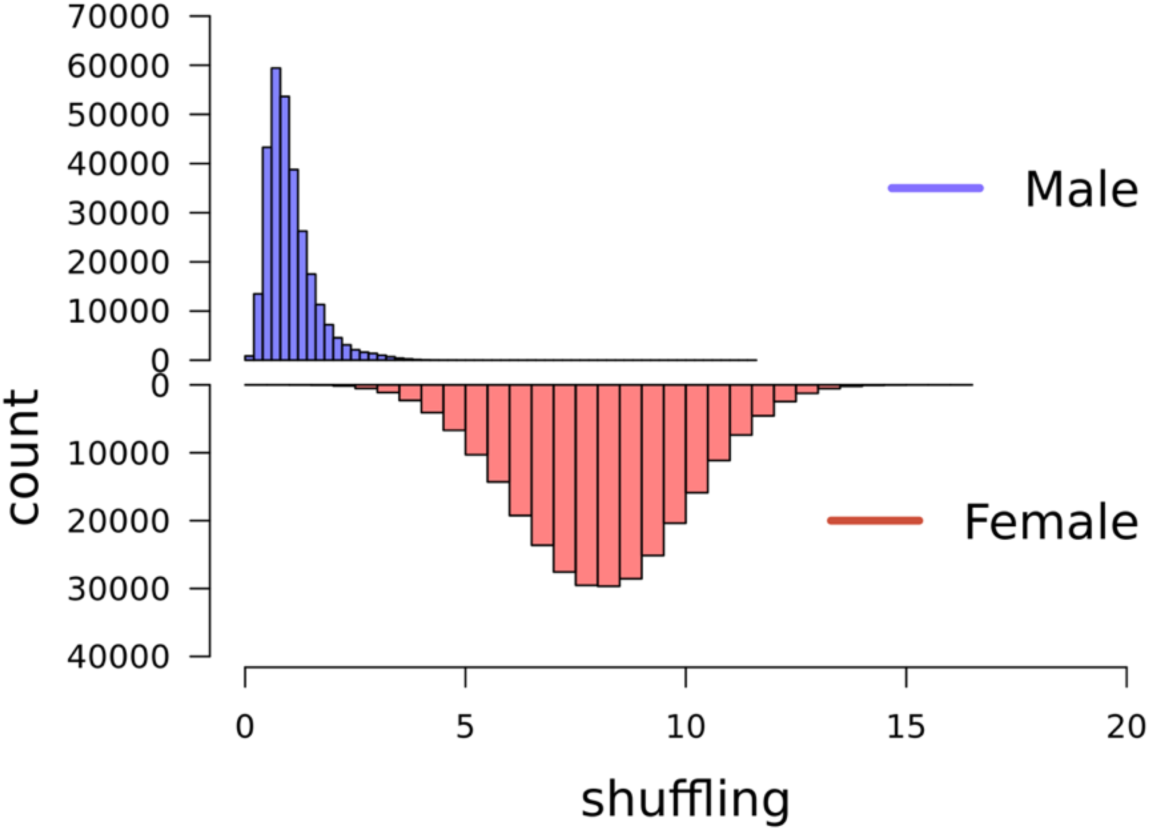
Distribution of intrachromosomal genetic shuffling 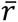. Maternal gametes in red and paternal gametes in blue.

**Table 2.** Results from variance component estimation of CC and 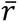.

| Trait | sex | N<sub>FIDs</sub> | N<sub>obs</sub> | mean (sd) | $h^2$(SE) | V<sub>p</sub> | V<sub>e</sub> |
| --- | --- | --- | --- | --- | --- | --- | --- |
| CC | female | 1580 | 287 063 | 19.6 (3.7) | 0.11 (0.01) | 14.2 | 11.7 |
| CC | male | 889 | 287 111 | 12.1 (2.9) | 0.10 (0.02) | 8.9 | 6.9 |
| $\bar{r}$ | female | 1580 | 287 063 | 8.06 (1.87) $\times 10^{-3}$ | 0.06 (0.01) | 3.8 $\times 10^{-3}$ | 3.1 $\times 10^{-3}$ |
| $\bar{r}$ | male | 889 | 287 111 | 1.01 (0.56) $\times 10^{-3}$ | 0.11 (0.03) | 0.4 $\times 10^{-3}$ | 0.2 $\times 10^{-3}$ |

N<sub>FIDs</sub> are the total number of FIDs (with repeated observations), N<sub>obs</sub> is the total number of observations (meiosis) for each sex. Mean is the mean CC or $\bar{r}$ with standard deviations in parenthesis. $h^2$ is the heritability estimate with standard errors in parenthesis. Effect f is the estimated effect of inbreeding (f) for each trait and sex.

### Genetic variation in measures of recombination

The heritability (h^2^) for CC was 0.11 (SE = 0.01) in females and 0.10 (SE = 0.02) in males. Genetic shuffling (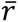) was also significantly heritable in both sex, but lower in females (h^2^ = 0.06, SE = 0.01) than in males (h^2^ = 0.11, SE = 0.03). Results from the variance component estimations for both traits is presented in table 2. The genetic correlations between CC and 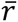 was 0.86 (0.01) in females, but only 0.42 (0.05) in males. The corresponding phenotypic correlations were 0.68 (0.00) for females and 0.43 (0.00) for females, i.e. phenotypic correlation between CC and 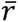 was lower than the genetic correlations for both sex. The genetic correlations between male and female CC and 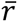 were 0.10 (0.06) and -0.10 (0.06), respectively (Table 3).

**Table 3.** Genetic and phenotypic correlations.

|  | **Male shuffling** | **Female shuffling** | **Male CC** | **Female CC** |
| --- | --- | --- | --- | --- |
| **Male shuffling** |  | - | 0.426<br>(0.003) | - |
| **Female shuffling** | -0.104<br>(0.060) |  | - | 0.679<br>(0.002) |
| **Male CC** | 0.646<br>(0.022) | - |  | - |
| **Female CC** | - | 0.892<br>(0.008) | 0.102<br>(0.060) |  |

In the lower triangle are genetic correlations between males and females for both traits with standard errors in parenthesis. Phenotypic Pearson's correlations between the two traits for each sex is in the upper triangle.

### Genome wide association studies

Genome wide association studies did not identify any significant quantitative trait loci for either crossover count or shuffling in males or females. In the GWA analysis for female CC and shuffling, there were one and three markers, respectively, that reached the significance threshold, but they were not supported by other markers in the same region and there were no clear candidate genes close to any of these markers (Fig 5 and 6). The top ten SNP markers for each GWAS are reported in Supplementary Table 1.

**Fig 5.**
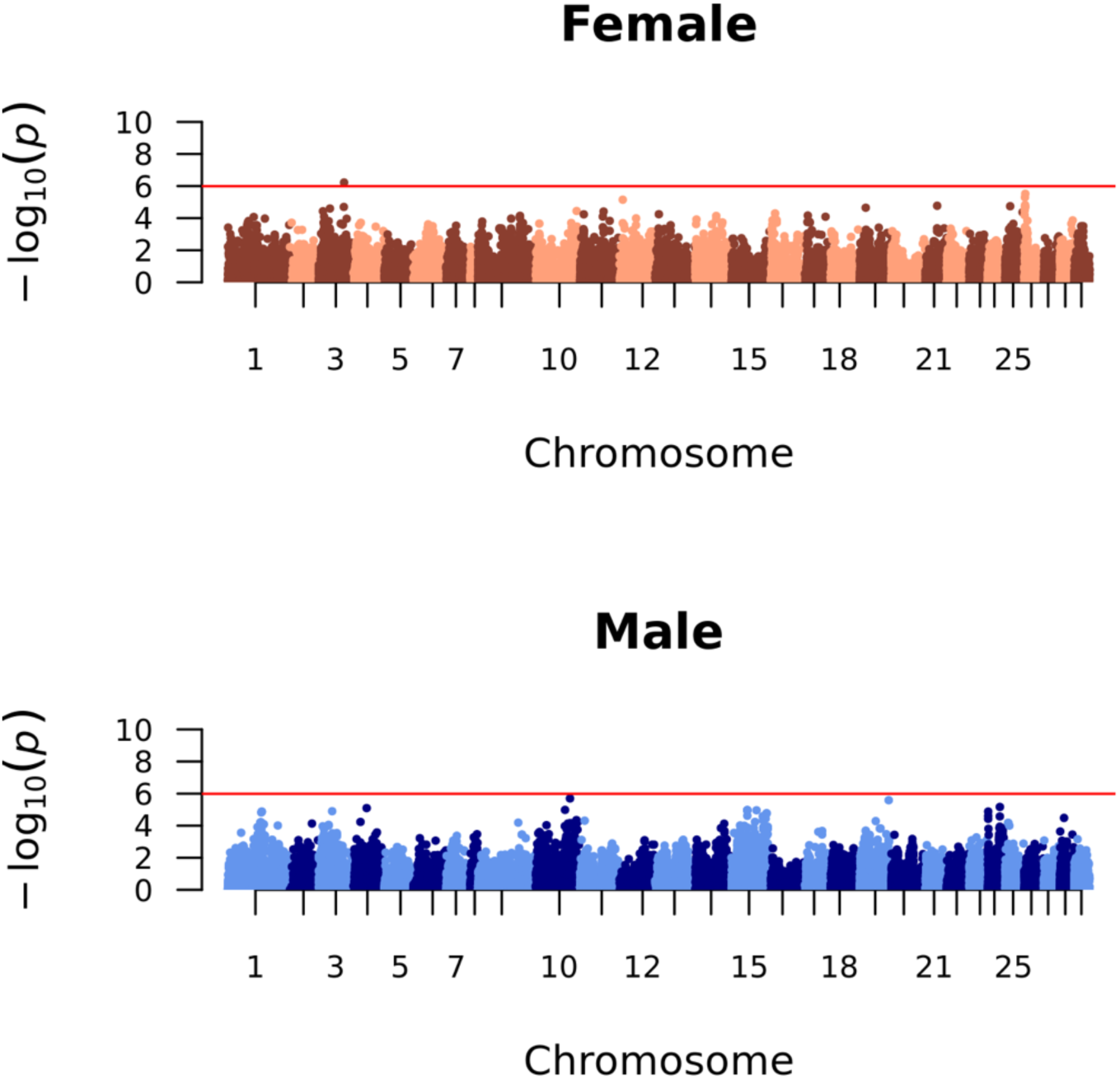
Manhattan plots of genome-wide association between markers and crossover count (CC). The red line is the genome-wide significance threshold using a Bonferroni correction. The Y axis is the negative logarithm of the p-values, and the x-axis is the physical positions of the markers with alternating colours for chromosomes 1-29 for males (A) and females (B).

**Fig 6.**
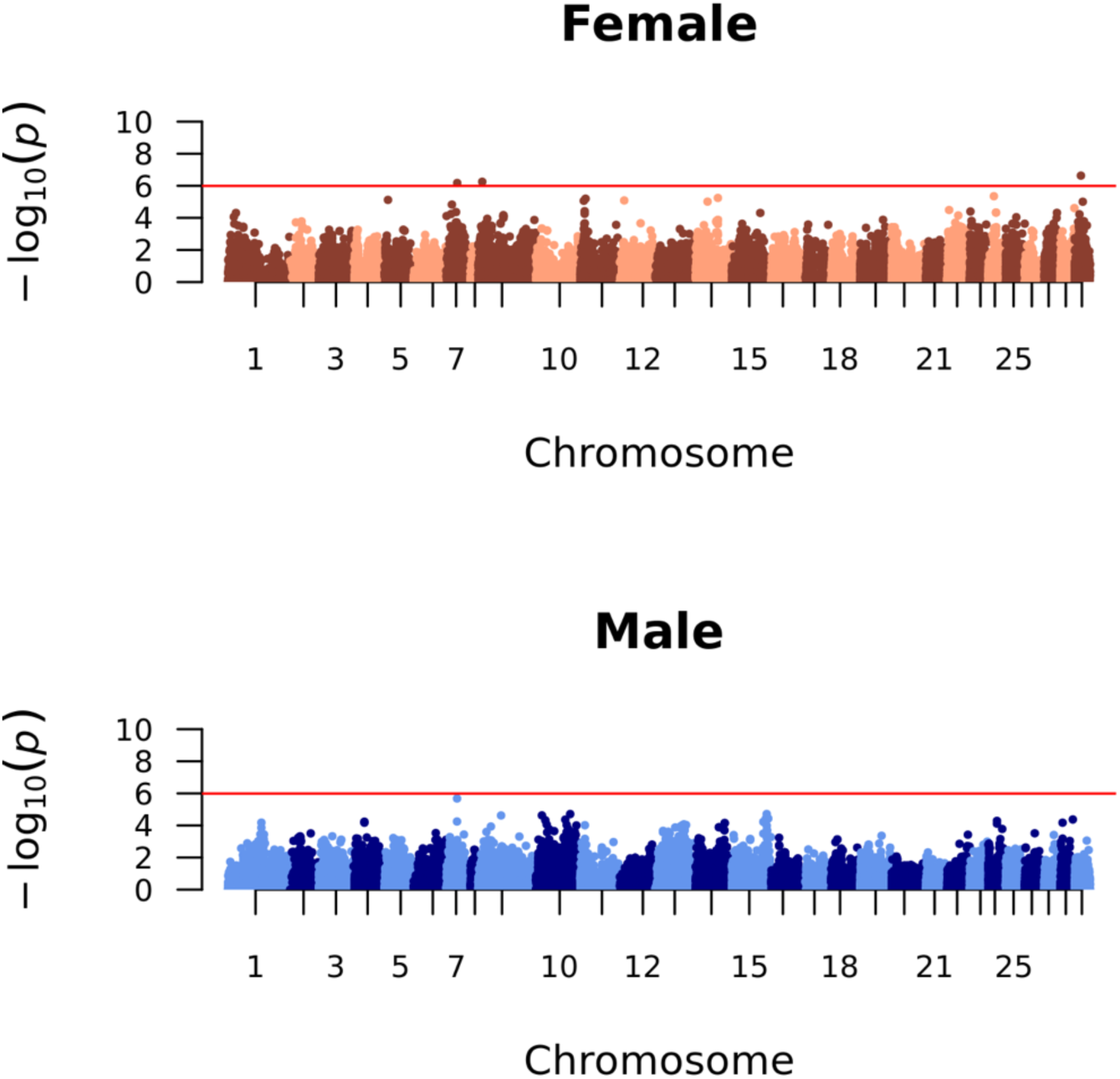
Manhattan plots of genome-wide association between markers and intrachromosomal shuffling (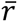). The red line is the genome-wide significance threshold = 0.05/Number of markers per analysis. The Y axis is the negative logarithm of the p-values, and the x-axis is the physical positions of the markers with alternating colours for chromosomes 1-29 for males (A) and females (B).

## Discussion

In this study, we confirm substantial sex-differences in genome-wide recombination rates and landscapes in Atlantic salmon, as previously reported by Lien et al (42) and Gonen et al. (43). Our study is the first to investigate this variation at the individual level, both in terms of crossover count (CC) and the degree to which alleles are shuffled on chromosomes to create new haplotypes (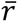). We showed that CC is heritable in both males and females, but with a low inter-sex genetic correlation, suggesting that the genetic architecture of CC is sex specific. Similarly, 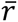 is heritable in females, but not in males. Our findings show that whilst females have ∼1.6 times more crossovers than males, they have eight times more genetic shuffling of alleles from one generation to the next, due to the extreme sex differences in crossover locations. We do not find any quantitative trait loci for either trait or sex, suggesting that the genetic variation in the two measures of recombination is polygenic. Here, we explore the results in more detail and discuss how the findings may be relevant in the breeding work on Atlantic Salmon, as well as how they contribute to the understanding of variation in rates and patterns of recombination in general.

### Extreme sex-differences in crossover landscapes

The sex-specific linkage map showed a female-biased heterochiasmy in salmon, but more striking was the large differences in the distribution of crossovers along the chromosomes. Male crossovers occured strictly in the telomeric regions, while female crossovers tend to occur closer to the centromeres, leaving regions in the middle of the acrocentric chromosomes (ssa8 to ssa29) with very low rates in both males and females (Fig 2). The total length in cM of the male and female linkage maps was shorter than previously published maps (42,43). This may be due to more accurate inference of physical marker positions relative to the assembled genome, rather than *de novo* inference of marker loci based on genetic linkage; incorrectly placed markers can often lead to inflation in map lengths. Also, the number of families and individuals used to create the linkage maps in this study is much higher than in previous published maps, with our full-sib family structures well suited for linkage mapping. Nevertheless, the relative difference between the sexes was similar to both studies, with females showing ∼1.38 and ∼ 1.5 times longer maps than males in Lien et al. (2011) and Gonen et al. (2014), respectively.

On the chromosomes with the largest sex differences in total genetic length (ssa2, ssa8 and ssa17), the male maps had little to no recombination in one or both telomeric regions of the chromosome (Fig 1). These regions coincide with regions reported in the paper by Lien et al (45) to have blocks of > 90% sequence similarity with blocks on other chromosomes.

Therefore, a potential explanation for the lack of crossovers between homologs in these regions is that these chromosomes are experiencing delayed rediploidization and are forming quadrivalents during meiosis (45). Cytological studies in different salmonid species find that multivalent pairing happens between the chromosomes with high sequence similarity and that the phenomena occur primarily in males (39,41). The high sequence similarity and tetrasomic inheritance makes these areas difficult to map, and these regions are indeed characterised by low marker density in our dataset. This means that there might be crossovers occurring in these regions that we are unable to pick up. In addition, if there is homeologue pairing in these regions, recombination events occurring between the two homeologue chromosomes may lead to an underestimation of genome-wide recombination rates in male Atlantic salmon; removing these three chromosomes reduces the sex difference in map lengths from 1.47 times higher in females to 1.33 times higher. Mechanistic explanations for the extreme difference in crossover distribution in males and females remain unresolved. One compelling avenue for further study in Salmonids, is that in other autotetraploid species such as *Arabidopsis arenosa*, recombination rates are reduced, and crossover interference appears to increase (46–48). Therefore, increased homeologue pairing in males may benefit from reduced recombination combined with telomeric crossing-over in order to mitigate issues arising from quadrivalent formation and/or mispairing of chromosomes during the crossover process.

Considering the evolutionary consequences of our findings, male recombination at the very end of the chromosomes leaves the haplotypes almost intact, which may be beneficial in a successful male to preserve their advantageous allelic combinations, particularly if males experience stronger diploid and/or haploid selection (28,31). However, the same could be argued for other species that do not share the same recombination patterns, yet may have stronger differences in selection between the sexes (e.g. with stronger sexual dimorphism and/or differential investment in gametes and offspring; (33,49). However, we cannot rule out similar importance of purely mechanistic suggestions, such as difference in timing of meiosis, chromatin structure and synaptonemal complex length between males and females resulting in sex-specific regions accessible for the recombination machinery at the time of meiosis or differences in the strength of crossover interference (49,50).

### Genetic variation in individual measures of recombination

To the best of our knowledge, these are the first results on genetic variation for individual measures of recombination in Atlantic salmon. Heritability for CC is moderate, 0.10 in males and 0.11 in females. This is consistent with estimates in other vertebrate species, which range from around 0.05 – 0.18 in pigs (16,51), sheep (12,13), cattle (15,21,52) and red deer (20), to as high as 0.41 and 0.46 in some *Drosophila* strains (17) and mouse lines (10), respectively. The genetic correlation between male and female CC in our study was very low at 0.10 (0.06). Despite the relatively large confidence interval, this estimate indicates that different loci affect CC in males and females. Intra-chromosomal shuffling 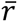 was also significantly heritable in females (h^2^ = 0.06) and males (h^2^ = 0.11). Considering the genetic correlation between CC and 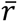 within each sex, we showed that this correlation was substantially lower in males than in females, 0.43 and 0.68 respectively.

We did not find any QTLs in either sex for either trait in the genome-wide association analysis, suggesting that these traits are polygenic in Atlantic salmon. This is in contrast to studies of individual recombination rates in mammals, where most studies point to a conserved oligogenic architecture of the trait (13,14,16,22). However, we cannot rule out that our dataset has reduced power to detect loci with a modest effect on recombination.

Compared to other studies in livestock (51) and human (27), the number of FIDs in our study are lower, impacting our ability to detect significant trait loci.

### What are the implications for breeding?

Our results suggest that there is the genetic potential to increase or decrease the amount of allelic shuffling and/or crossover counts within Atlantic salmon, and that this can be achieved independently in males and females, due to low genetic correlations between the sexes. However, the relatively low heritabilities and polygenic nature of these traits suggest that genetic change may be slow because the many small effect loci can be pleiotropic and/or linked with genes affecting other traits prioritised in selection, and that a corresponding phenotypic change will be modest (53). Previous focus on the relevance of recombination for animal breeding has been local or genome-wide rates of crossovers (53–55). In our study, the striking difference between male and female shuffling in Atlantic salmon demonstrates the critical importance of crossover for shuffling and creation of novel haplotypes that may have previously been overlooked. There are considerable differences in the probability of shuffling between linked loci in male and female meiosis, with males transmitting haplotypes to offspring that are relatively unchanged. However, we must also consider that each offspring will always inherit exactly one paternal and one maternal gamete and in the subsequent generations, alleles will segregate independently of which sex they were transmitted from in previous generations, so the population level effect of the conserved haplotypes in male salmon may be limited. Furthermore, little is known about the potential biological and fitness consequences of altering rates or distribution of crossovers. The mechanisms leading to the observed variation in shuffling are likely associated with the mechanisms that control crossover location and rate. For example, if low rates of crossovers in both males and females in chromosome centres is associated with chromosome fusions following the whole genome duplication, attempting to select for crossover location closer to chromosome centres may cause reduction in fertility (56). Similarly, despite some evidence for a potential to increase genetic gain with higher recombination rates (53), selecting for higher recombination rates is not likely to be beneficial or even possible with crossover interference. The negative consequences of extensive recombination rates remain to be understood, although higher mutagenic load is likely (14), combined with the fact that there seems to be an upper limit for number of crossovers per chromosome shared among species across a broad selection of taxa (8),

### Conclusions and future directions

In conclusion, this study shows that there is genetic variation in genome-wide rates and distribution of recombination in Atlantic salmon. Consistent with previous studies, Atlantic salmon show extreme levels of heterochiasmy, especially in the distribution of crossover events along the chromosomes. We show that the genetic correlation between male and female rates and distribution of crossovers is very low, suggesting that they behave and may be altered as different traits in males and females. Future studies should aim to include the multivalent pairing in male meiosis to get the full picture of male crossover rates and distribution. Overall, these findings provide a basis to better understand the causes and consequences of recombination rate variation in general, and the genetic architecture of the trait in Atlantic salmon specifically.

## Methods

### Atlantic salmon genetic dataset

Genotypes from a total of 375,381 individuals were available from the Norwegian AquaGen Atlantic Salmon (*Salmo salar*) breeding population for this study. The breeding work in this population started in 1970 and it has founders from 41 Norwegian rivers. The individuals in this dataset were born between year 2008 and 2020. Individuals were genotyped on two different customized SNP arrays, Affymetrix custom 49K and Affymetrix custom 70K. Genotype calls were generated using the Thermo Fisher Best Practices Genotyping Analysis Workflow. Markers in the categories PolyHighResolution and NoMinorHom were kept for further analysis, i.e SNPs with well-separated genotype clusters and two or more alleles in the genotype calls, and where one cluster is homozygous, and one is heterozygous for biallelic SNPs and only one homozygous cluster and one or more heterozygous cluster appear for multiallelic SNPs (57). Finally, only markers common to both SNP arrays were kept for further analysis. resulting in a set of 35,033 markers and a total genotyping call rate of 0.997 in the final dataset. The physical positions of the SNPs were determined based on the Atlantic salmon reference genome assembly Ssal_v3.1 (GenBank accession GCA_905237065.1). This data set is referred to hereafter as the 35K dataset. All focal individuals (see section three generation full-sib families below) were also imputed to the 49.087 SNP markers on the customized Affymetrix 49K chip; this dataset was used for the genome wide association analysis only.

### Three generation full-sib families

The pedigree was ordered into full-sib families as follows: for each unique sire – dam mating pair, hereafter referred to as the focal individuals or FIDs, we constructed a three-generation family that included their offspring and potentially genotyped parents (see Fig 7 for illustration of the family structure). This enables phasing of the gametes transmitted from the FIDs to the offspring, in turn identifying the crossover positions that occurred during meiosis in the FIDs. Therefore, recombination phenotypes (i.e. crossover count and intra-chromosomal shuffling) are assigned to the FIDs. An FID can be in several families if the individual is mated with several individuals in the pedigree, or as an offspring or grandparent, but our study design means that each meiosis was only counted once. The 35K set had a total of 5568 unique full-sib families with number of offspring ranging from 1 to 537. Because the number of full-sibs were high in most of the families, genotyped grandparents were not vital for proper phasing of the offspring gametes and therefore not set as a strict criteria for inclusion.

**Fig 7.**
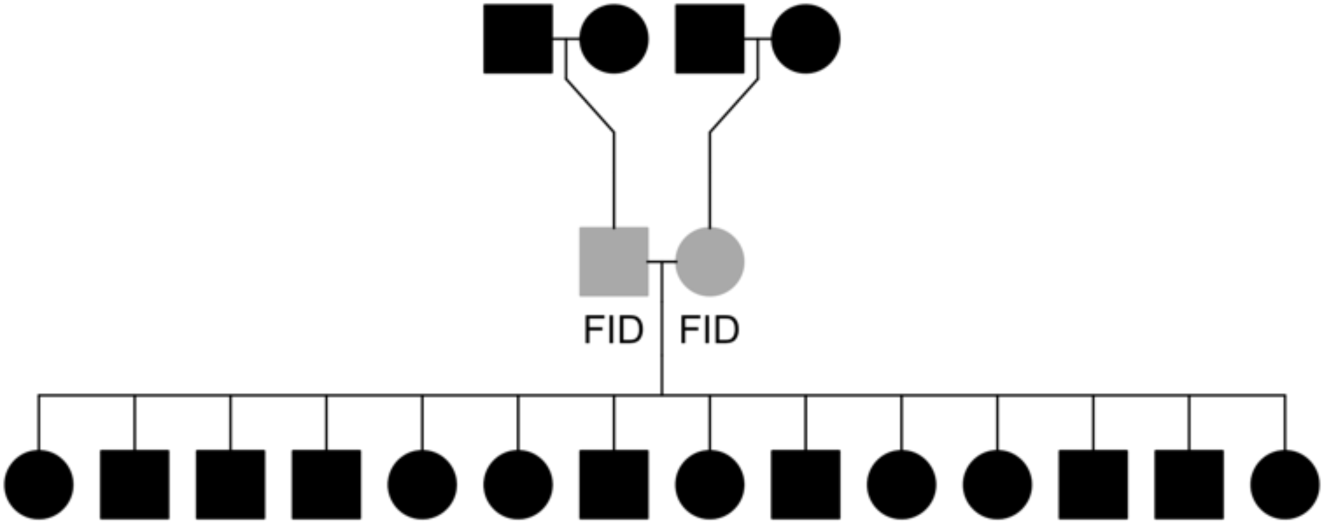
Illustration of the full sib family structures. This structure is used to phase gametes that are transmitted from focal individuals (FIDs, in grey) to their offspring. In cases where parents in the FIDs are known, they are included to improve phasing accuracy.

### Linkage mapping

Linkage mapping was conducted in LepMap3 (58) assuming marker orders were the same as on the Atlantic salmon genome and that each chromosome constituted a linkage group. The *filtering2* module was run as suggested for multi-family datasets with a *datatolerance* = 0.01 to filter markers based on segregation distortion. The *separatechromosomes2* module was run within linkage group and markers that were not assigned to the main group (LOD score < 5) were excluded, as suggested in species where chromosome-level assemblies and marker positions are well established. The *ordermarkers2* module was run with the option to evaluate the given marker order, i.e. to calculate the centimorgan (cM) positions using the Haldane mapping function option. A total of 35,032 markers across 29 chromosomes were included in the final linkage map. These data were then used to calculate approximate fine-scale recombination rates across the genome. SNPs were assigned to bins of 1Mb based on their genomic positions, and the cM/Mb rate was measured as the difference in cM divided by the difference in Mb between the first and last SNP marker within each bin. Finally, it should be noted that Atlantic salmon do not have distinct sex chromosomes but have an autosomal sex-determining region on Ssa3 (59).

### Estimation of individual recombination rates

Crossover count (CC) was determined for each gamete transmitted from an FID to their offspring from gamete-phased output of the *orderMarkers2* module in LepMap3, and assigned to the FID in which the meiosis took place. Intra-chromosomal genetic shuffling, 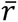, was calculated as the probability that a randomly chosen pair of loci on the same chromosome was unpaired during gamete production (in meiosis) following the method suggested by Veller et al. (37):

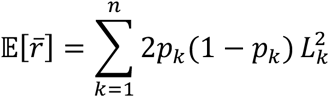

where *k* is the chromosome number 1-29, *p* is the proportion of paternally inherited alleles, *1-p* is the proportion of maternally inherited alleles, and *L* is the length of the chromosome as a fraction of the total length of the genome. This method determines 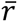 for each gamete and is assigned as a phenotype of the FID in which meiosis took place.

### Genetic variation in measures of recombination

Variance components for individual CC and 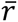 were estimated in DMU v6 (60) with a repeatability model using the Restricted Maximum Likelihood (REML) and average information (AI) algorithm. The model was:

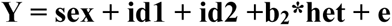

where Y is the response variable (CC or 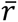), **sex** is the fixed effect of sex, **id1** is the random additive genetic effect of the FID (N= 2,469) with a covariance matrix proportional to the genomic relationship matrix, **id2** is the random effect of the FIDs permanent environment (i.e. environmental effects that are constant across repeated measures on an FID), **het** is the individual inbreeding coefficient (method-of-moments F) calculated with the *--het* function in PLINK v1.9 (61), **b_2_** is the regression of CC or 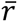 on **het** of the FID, and **e** is the residual effect. The narrow-sense heritability (*h^2^*) was defined as the proportion of phenotypic variance explained by the additive genetic effect **id1** (i.e. the estimated additive genetic variance divided by the sum of variances estimated for all random effects) and was estimated separately for each sex. Repeatability was measured as the sum of the genetic variance and permanent environment variance divided by the sum of all variances estimated for all random effects.

### Genome wide association studies (GWAS)

We conducted GWAS for CC and 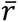 using the 49K SNP dataset using a mixed linear model-based analysis in GCTA version 1.93.2a_beta (62), where the chromosome that the focal SNP is located on is left out of the genomic relationship matrix as implemented in the *--mlma-loco* module. The model was structured as follows:

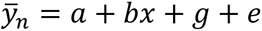

Where 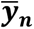 is the mean CC or 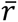 for *n* observations on the individual, *a* is the trait mean, *b* is the fixed additive effect of the SNP tested for association and *x* is the SNP genotypes 0, 1 and 2 for the homozygote, heterozygote and opposite homozygote respectively, *g* is a vector of random effects assumed to 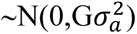 where G is the genomic relatedness matrix calculated with SNP markers on all the autosomes except the chromosome of the SNP currently tested for association and 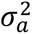 is the genetic variance, and *e* is the residual term 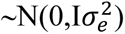. To account for *n* multiple observations per FID, 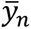 was weighted based on the number of observations, and trait and sex-specific heritabilities (h^2^) and repeatabilities (t) estimated following the method from Garrick et al., (63). The significance threshold for association was determined using a Bonferroni correction at α = 0.05, determined as P = 0.05/49,087 = 1.02 ×10^-6^.

## Acknowledgments

We want to thank Aquagen for providing the data for this study. Special thanks to Pasi Rastas who gave advice about the lepmap3 software used for linkage mapping. We thank Sigbjørn Lien, Nicola Barson, Arne B. Gjuvsland and John McAuley for helpful discussions. Most of the analyses in this study have been executed on the Orion cluster at NMBU (https://orion.nmbu.no).

## Author contribution

**Conceptualization:** PB, SEJ, CB

**Data curation:** CB, TMK

**Formal analysis:** CB

**Writing – original draft:** CB

**Writing – review and editing:** SEJ, CB, TMK, PB.

**Visualization:** CB

**Funding acquisition:** PB

**Investigation:** CB, SEJ, TMK, PB

**Methodology:** SEJ, CB, TMK

**Project administration:** PB

**Supervision:** PB, SEJ

---

**Table S1.** Linkage map. Male and female genetic positions in cM for all SNP markers in the 35K dataset. See additional file.

**Table S2.** Top 10 SNPs from GWAS on shuffling and crossover count in both sex.

| Trait | Sex | Chr | SNP name | bp | Freq | b | SE | p |
| --- | --- | --- | --- | --- | --- | --- | --- | --- |
| CC | Male | 10 | Affx-87165681 | 97158781 | 0.216 | 0.936 | 0.197 | 2.00E-06 |
|  |  | 19 | Affx-986688163 | 80841039 | 0.209 | -0.844 | 0.179 | 2.58E-06 |
|  |  | 24 | Affx-517031085 | 33571077 | 0.286 | 0.778 | 0.173 | 6.79E-06 |
|  |  | 4 | Affx-87621822 | 40639301 | 0.177 | 0.890 | 0.199 | 8.01E-06 |
|  |  | 10 | Affx-86913301 | 83219166 | 0.176 | 0.850 | 0.193 | 1.04E-05 |
|  |  | 15 | Affx-87001560 | 44530526 | 0.517 | 0.651 | 0.148 | 1.09E-05 |
|  |  | 15 | Affx-87481996 | 44534054 | 0.517 | 0.651 | 0.148 | 1.09E-05 |
|  |  | 15 | Affx-87658641 | 44541415 | 0.483 | -0.651 | 0.148 | 1.09E-05 |
|  |  | 15 | Affx-98990447 | 44543717 | 0.483 | -0.651 | 0.148 | 1.09E-05 |
|  |  | 15 | Affx-517023952 | 70320171 | 0.407 | -0.656 | 0.149 | 1.10E-05 |
| CC | Female | 3 | Affx-87260986 | 71736893 | 0.286 | -0.643 | 0.129 | 6.00E-07 |
|  |  | 26 | Affx-517021201 | 3373590 | 0.327 | -0.567 | 0.121 | 3.01E-06 |
|  |  | 26 | Affx-87324563 | 2403259 | 0.221 | 0.662 | 0.145 | 5.06E-06 |
|  |  | 12 | Affx-93396222 | 10860972 | 0.349 | -0.538 | 0.120 | 7.09E-06 |
|  |  | 26 | Affx-86915404 | 2971226 | 0.450 | 0.498 | 0.115 | 1.52E-05 |
|  |  | 21 | Affx-86977045 | 32739741 | 0.192 | -0.626 | 0.146 | 1.69E-05 |
|  |  | 25 | Affx-86916185 | 14361093 | 0.181 | 0.628 | 0.147 | 1.81E-05 |
|  |  | 26 | Affx-87723009 | 2274470 | 0.285 | 0.582 | 0.136 | 1.85E-05 |
|  |  | 26 | Affx-87775881 | 2569233 | 0.267 | -0.552 | 0.129 | 1.91E-05 |
|  |  | 3 | Affx-87597243 | 71065500 | 0.498 | -0.496 | 0.116 | 2.01E-05 |
| Shuffling | Male | 7 | Affx-87702070 | 30082074 | 0.177 | 0.253 | 0.053 | 2.08E-06 |
|  |  | 15 | Affx-87438278 | 97920722 | 0.201 | 0.208 | 0.049 | 1.94E-05 |
|  |  | 10 | Affx-87814965 | 96995476 | 0.399 | 0.180 | 0.042 | 1.94E-05 |
|  |  | 10 | Affx-87169893 | 17900600 | 0.567 | -0.179 | 0.042 | 2.34E-05 |
|  |  | 9 | Affx-87137995 | 63788953 | 0.370 | 0.197 | 0.047 | 2.37E-05 |
|  |  | 15 | Affx-87191200 | 95892936 | 0.239 | 0.194 | 0.047 | 3.65E-05 |
|  |  | 15 | Affx-87285851 | 101670539 | 0.192 | 0.215 | 0.052 | 3.70E-05 |
|  |  | 15 | Affx-87285853 | 101671231 | 0.192 | 0.215 | 0.052 | 3.70E-05 |
|  |  | 28 | Affx-87026689 | 35657626 | 0.198 | 0.218 | 0.053 | 4.24E-05 |
|  |  | 15 | Affx-87320647 | 97937595 | 0.315 | 0.176 | 0.043 | 4.27E-05 |
| Shuffling | Female | 29 | Affx-87275877 | 17198796 | 0.200 | 0.252 | 0.049 | 2.32E-07 |
|  |  | 9 | Affx-87805131 | 11090861 | 0.539 | -0.198 | 0.040 | 5.58E-07 |
|  |  | 7 | Affx-87197793 | 30511902 | 0.196 | -0.256 | 0.051 | 6.68E-07 |
|  |  | 24 | Affx-87658020 | 15513356 | 0.339 | -0.182 | 0.040 | 4.46E-06 |
|  |  | 14 | Affx-87347934 | 62926154 | 0.440 | -0.175 | 0.039 | 5.71E-06 |
|  |  | 11 | Affx-87566793 | 14623343 | 0.219 | 0.209 | 0.046 | 6.33E-06 |
|  |  | 5 | Affx-87225703 | 10195400 | 0.290 | 0.193 | 0.043 | 7.59E-06 |
|  |  | 12 | Affx-87008423 | 14255690 | 0.218 | 0.214 | 0.048 | 8.24E-06 |
|  |  | 11 | Affx-86997783 | 9768960 | 0.491 | 0.178 | 0.040 | 8.65E-06 |
|  |  | 14 | Affx-87808468 | 33998757 | 0.132 | -0.242 | 0.055 | 9.73E-06 |

**bp** is the basepair position of the SNP marker, **Freq** is the allele frequency of the effect allele, **b** is the the effect estimate and **SE** the subsequent standard error of the effect estimate, **p** is the p value.

